# Mouse Cortical Cellular Diversification Through Lineage Progression of Radial Glia

**DOI:** 10.1101/2025.01.24.634707

**Authors:** Lin Yang, Ziwu Wang, Yanjing Gao, Zhenmeiyu Li, Guoping Liu, Zhejun Xu, Zhuangzhi Zhang, Yan You, Zhengang Yang, Xiaosu Li

## Abstract

Cortical radial glia (RGs) sequentially generate pyramidal neurons (PyNs) and glia. In this study, we investigated the cell intrinsic programs underlying cortical cellular diversification using time-series scRNA-seq and snATAC-seq on purified mouse cortical progenitors across embryonic and postnatal stages. Our data revealed that RGs transition from early to late over time, sequentially producing intermediate neuronal progenitors (INPs) and intermediate glial progenitors (IGPs). While INPs expand exclusively to generate PyNs, IGPs progress from young to old, sequentially producing cortical astrocytes, oligodendrocytes, and olfactory bulb interneurons. We constructed comprehensive molecular maps that reflect cell lineage progression. In particularly, we found that chromatin accessibility drives cortical cellular diversification by restricting broadly expressed transcription factors to specific stages and cell types. Developmental changes in chromatin accessibility confine *Lhx2*-induced neurogenesis to early-stage RGs, leading to the loss of neurogenic competence and the acquisition of gliogenic competence as corticogenesis progresses.

## Introduction

During mammalian corticogenesis, cortical radial glia (RGs) act as neural stem cells (NSCs), sequentially producing cortical pyramidal neurons (PyNs) and glial cells (Kriegstein and Alvarez-Buylla 2009). Cortical RGs integrate multiple extrinsic molecular cues to regulate the timing of the neurogenesis-to-gliogenesis transition (Miller and Gauthier 2007). Several gliogenic induction cues, such as JAK-STAT and BMP, efficiently promote gliogenesis in late-stage RGs, but not in early-stage RGs (He et al. 2005; Li et al.; Gross et al.). Early- and late-stage RGs respond differently to gliogenic cues, suggesting that intrinsic changes in RGs determine their cellular competence to respond, or not, to gliogenic induction cues. Developmental DNA demethylation at the promoters of gliogenic genes is coordinated with the neurogenesis-to-gliogenesis transition. The promoters of *Gfap* and *S100b* are hypermethylated during the early stages of corticogenesis (Takizawa et al. 2001a). Notch activation disrupts the interaction between the *Gfap* promoter and the maintenance methyltransferase Dnmt1, inducing demethylation of the *Gfap* promoter and promoting accessibility to STAT transcription factors (Namihira et al. 2009; Fan et al. 2005a; He et al. 2005; Takizawa et al. 2001a; Bonni et al. 1997). Despite DNA demethylation, other cell-intrinsic programs underlying the RG competence switch from neurogenic to gliogenic remain relatively unexplored.

To systematically investigate the cell intrinsic programs underlying RG competence switch from neurogenic to gliogenic, we performed Single cell RNA sequencing (scRNA-seq) and single nucleus ATAC sequencing (snATAC-seq) on purified cortical progenitors. scRNA-seq and snATAC-seq are powerful tools for identifying cell molecular profiles and epigenetic status. Several recent studies have employed scRNA-seq and/or snATAC-seq to investigate mouse cortical development (Telley et al. 2019; Ruan et al. 2021; Di Bella et al. 2021; La Manno et al. 2021). These studies have provided a comprehensive atlas of corticogenesis and the temporal patterning of early-stage cortical progenitors. However, as corticogenesis progresses to the later stages, the accumulated postmitotic PyNs become the predominant cell type, with progenitors and glial cells accounting for less than 10% of all cortical cells. The neurogenesis-to-gliogenesis transition and the temporal patterning of late-stage progenitors have not been thoroughly explored. While early-stage cortical RGs exclusively produce PyNs, late-stage RGs simultaneously generate PyNs, astrocytes, oligodendrocytes, and OB interneurons (Li et al. 2021; Yang et al. 2022; Merkle et al. 2007). To elucidate the molecular logic underlying the neurogenesis-to-gliogenesis transition and the complex lineage diversification during late corticogenesis, it is essential to use purified cortical progenitors and glial lineage cells for scRNA-seq and snATAC-seq.

In this study, we performed time series scRNA-seq and snATAC-seq on FlashTag (FT) and *hGFAP-GFP* labeled cortical progenitors. During the neurogenic stages (before E16), RGs exclusively generated PyNs, either directly or indirectly, via intermediate neuronal progenitors (INPs). As corticogenesis progresses to the gliogenic stages, subpopulations of RGs transition to gliogenesis. Gliogenic RGs generated a common population of intermediate glial progenitors (IGPs), which transition from young to old over time, sequentially generating cortical astrocytes, oligodendrocytes, and olfactory bulb (OB) interneurons. We established comprehensive molecular maps for cortical lineage commitment and cellular diversification, linking gene modules to specific cell fates and differentiation stages, with molecular developmental trajectories that mirror cell lineage trajectories. The transcriptome and epigenome in cortical RGs were temporally dynamic, coordinating the temporal neurogenesis-to-gliogenesis switch. In particularly, chromatin accessibility drives cortical cellular diversification by restricting broadly expressed transcription factors to specific stages and cell types. Developmental changes in chromatin accessibility confine *Lhx2*-induced neurogenesis to early-stage RGs. As RGs progress to late stages, their responsiveness to Lhx2 diminishes due to decreased chromatin accessibility at Lhx2 target genes, thereby facilitating the transition from neurogenesis to gliogenesis.

## Results

### Two Axes of Temporal Patterning Govern Radial Glia Lineage Progression

To elucidate the temporal patterning mechanisms of mouse cortical development, we performed time-series scRNA-seq on cortical progenitors from embryonic day 13 (E13) to postnatal day 1 (P1). We employed three strategies to isolate cortical progenitors (Fig. 1A). At E13, whole cortices were isolated and dissociated into single cells. From E15 to E18, cortical progenitors were enriched using fluorescence-activated cell sorting (FACS) following 24 hours of in utero intracerebroventricular FlashTag (FT) injection. From E18 to P1, GFP-expressing cells from *hGFAP-GFP* transgenic mouse cortices were sorted by FACS. The collected cells underwent single-cell 3’ gene expression library construction and high-throughput sequencing on an Illumina platform (Fig. 1A). Additionally, two E14 cortical samples from a previously published study were included (Noack et al. 2022). In total, 25276 cells from the neurogenic stages (E13-E15) and 40090 cells from the gliogenic stages (E17-P1) were recovered, representing the neurogenic and gliogenic datasets, respectively (Fig. 1D and Fig. S1A).

**Fig. 1:**
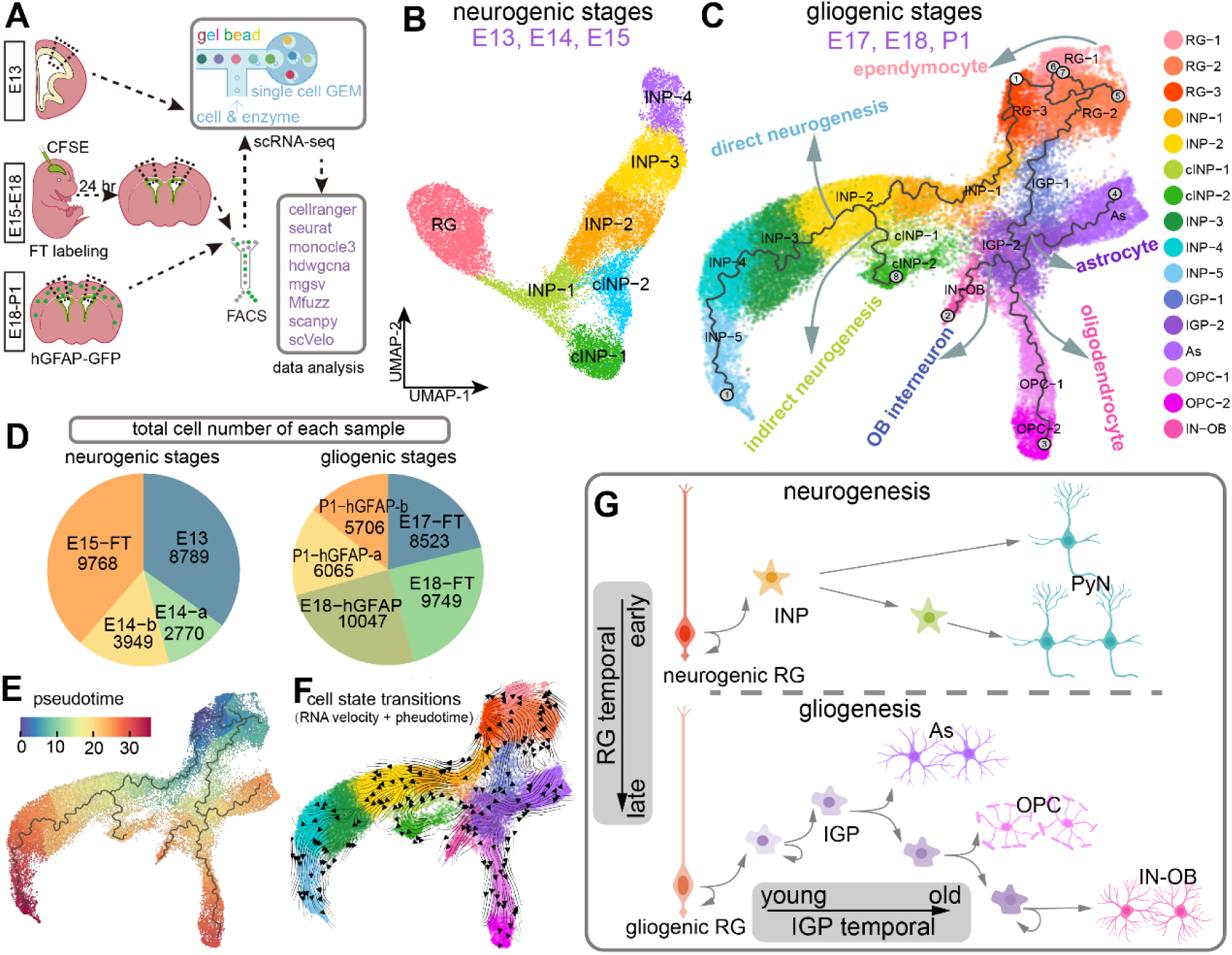
Time series scRNA-seq of the developing cortex. **A**: Experimental scheme for the time series scRNA-seq analysis of the developing cortex. **B-C**: UMAP projections of the combined scRNA-seq data from neurogenic stages (E13, E14 and E15) and from the gliogenic stages (E17, E18 and P1). The black lines represent cell lineage trajectories reconstructed using Monocle3, depicting the differentiation paths from root cells (circled number 1 with white background) to terminal cells (circled numbers with gray background) (C). RG, radial glia; INP, intermediate neuronal progenitor; cINP, cycling INP; IGP, intermediate glial progenitor; As, astrocyte; OPC, oligodendrocyte precursor cell; IN-OB, interneuron of olfactory bulb. **D**: Total cell number for each individual sample. **E**: UMAP visualization of the inferred pseudotime. **F**: Cell state transitions revealed through integration of RNA velocity analysis and Monocle pseudotime. **G**: Schematic diagram of cortical cellular diversification.

Cell type identities were determined based on the expression of well-known marker genes (Fig. 1B-C and Fig. S1B-C). During neurogenesis, *Pax6*+/*Sox2*+ cortical RGs exclusively produced *Eomes*+/*Neurog2*+ INPs. The INPs either directly differentiated into *Nueord6*+ PyNs (direct neurogenesis) or transformed into cycling INPs (cINPs), which underwent 1-2 rounds of proliferation before producing PyNs (indirect neurogenesis) (Fig. S1B) (Noctor et al. 2004). During gliogenesis, cell lineage trajectory analysis using monocle3 revealed that distinct RG subpopulations RG-3 and RG-2 produced INPs and *Olig2*+/*Egfr*+ intermediate glial progenitors (IGPs) for neurogenesis and gliogenesis, respectively (Fig. 1C). Consequently, RG-3 and RG-2 were classified as neurogenic and gliogenic RGs. Cell state transitions were estimated by integrating RNA velocity with pseudotime analysis, with results consistent with the lineage trajectories (Fig. 1F). Notably, the orientation of RNA velocity vectors from RG-3 to RG-2 suggested a transition from neurogenic to gliogenic RG states. While INPs exclusively differentiated into PyNs, either directly or indirectly, IGPs diversified into three distinct cell types: *Aldh1l1*+ astrocytes, *Sox10*+ oligodendrocytes, and *Dlx2*+ OB interneurons (Fig. S1C). Rather than generating separate progenitors for each cell type, RGs produced a common population of IGPs that transitioned from young to old in pseudotime, first producing astrocytes, followed by oligodendrocytes, and ultimately OB interneurons (Fig. 1C, E).

In summary, mouse cortical cellular diversification occurs through two axes of temporal patterning within RG lineages (Fig. 1G): first, RGs transition from early to late stages over time, sequentially producing INPs and IGPs; second, IGPs progress from young to old over pseudotime, sequentially generating cortical astrocytes, oligodendrocytes, and olfactory bulb interneurons.

### Dynamic Gene Expression during RG Lineage Progression

The transcriptional profiles of cortical RGs exhibit temporal dynamics throughout development. To systematically characterize these temporal expression patterns, we extracted RG-specific expression matrices from both time-series single-cell RNA sequencing (scRNA-seq) datasets and our previously published cortical scRNA-seq data (Li et al. 2021). A total of 17000 RGs were recovered from 14 samples, including eight timepoints spanning E13 to P2. After filtering out genes with low expression at any timepoints or with small variance, a total of 2656 genes were identified as RG temporal dynamic expression genes. To reveal the dynamic gene expression patterns, we applied soft clustering to these genes based on their expression levels using the fuzzy c-means algorithm. We identified 12 distinct clusters of temporal expression patterns (Fig. S2A-B). Clusters 3, 7, and 8 were downregulated over time (612 genes), while Clusters 6, 11, and 12 were upregulated (972 genes). The remaining six clusters exhibited oscillatory patterns (Fig. S2A-B). GO enrichment analysis revealed that temporally downregulated genes (early RG genes) were primarily involved in the cell cycle, RNA splicing, and chromatin remodeling. Conversely, temporally upregulated genes (late RG genes) were mainly involved in protein localization, cell junction organization, and gliogenesis (Fig. S2C). Notably, some of the temporally downregulated and upregulated genes were specifically expressed during the neurogenic and gliogenic stages, respectively (Fig. S2D).

Next, we examined the dynamic changes in the molecular program during glial lineage progression. Cells associated with the gliogenic trajectories—from the root cell to the three terminal endpoints corresponding to astrocytes, oligodendrocytes, and OB interneurons—were isolated and designated as the glial lineage (Fig. 2A). Genes dynamically expressed along lineage pseudotime were selected to construct a molecular map of glial lineage diversification. A total of 617 genes were clustered into seven distinct gene modules (Fig. 2B). Aggregated expression levels of individual modules showed that these modules were specifically expressed in distinct cell types (Fig. 2C). Each gene module was named after its corresponding cell types. The spatial adjacency between the IGP module and OL/IN-OB module was consistent with their cell lineage relationship. However, the As module was aberrantly positioned away from the IGP module, which conflicted with their lineage relationship. The RG & As module, which included genes expressed by both RGs and astrocytes, was positioned between the RG and As modules (Fig. 2B). Numerous genes shared between RGs and astrocytes may influence the spatial distribution of the gene modules, potentially explaining the inconsistency between cell lineage trajectories and gene module trajectories. Genes within the same module exhibited similar expression patterns, suggesting that they may work coordinately to contribute to the same developmental events.

**Fig. 2:**
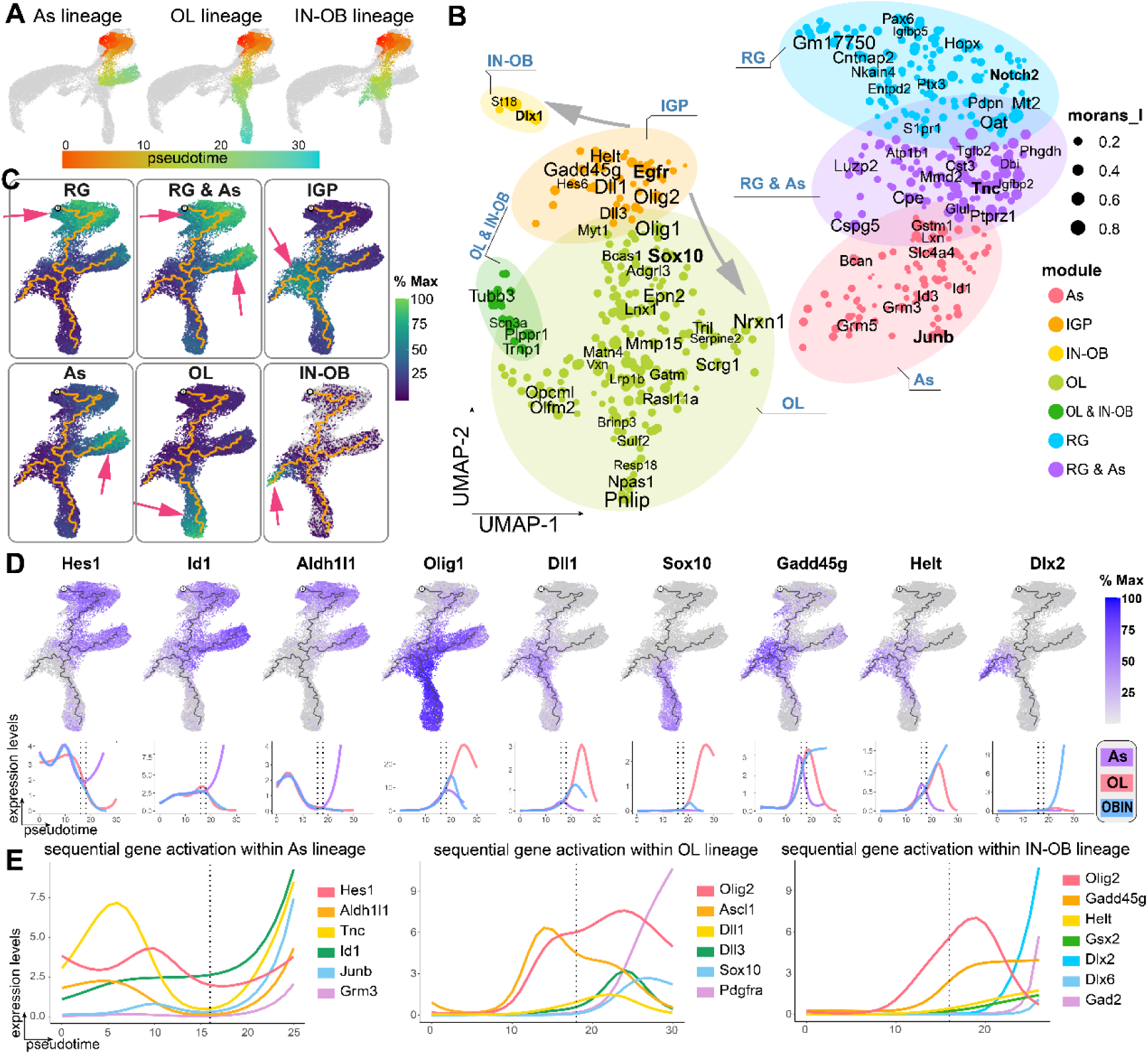
Molecular Maps Depicting Glia Lineage Diversification. **A**: UMAP visualization of each lineage cells, colored by pseudotime. **B**: A panoramic molecular map illustrating molecular dynamics along glia lineage progression, colored by gene modules. OL, oligodendrocyte. morans_I, spatial autocorrelation Moran’s index. **C**: UMAP visualization showing aggregated expression levels of each individual gene module. **D**: Gene expression dynamics along glia lineage progression. Upper panel: UMAP visualization of gene expression levels with lineage trajectories; lower panel: GAM smoothed gene expression levels along each lineage pseudotime, and the two vertical dashed lines mark the pseudotime corresponding to the branch points. **E:** Sequential gene activation patterns along each lineage pseudotime.

Hes1 and Id1 were expressed in young IGPs and cells along the astrocyte lineage trajectory (Fig. 2D, F). As the trajectory passed through the astrocyte bifurcation point, Hes1 and Id1 expression was extinguished in old IGPs. Subsequently, old IGPs activated Dll1 and Dll3 expression while increasing Olig1 and Olig2 expression, biasing them toward the oligodendrocyte lineage (Fig. 2D, F). Eventually, IGPs upregulated Gadd45g, Helt, and Gsx2 expression, committing to an OB interneuron fate (Fig. 2D, F).

In summary, cellular molecular profiles change along two axes of temporal lineage progression. First, RGs temporally express distinct gene sets, likely coordinating the transition from neurogenesis to gliogenesis. Second, IGPs sequentially express combinatorial gene sets, likely orchestrating the stepwise generation of cortical astrocytes, oligodendrocytes, and OB interneurons.

### *LHX2* Suppresses the Neurogenesis-to-Gliogenesis Transition in the Cortex

In *Drosophila*, the temporal patterning of NSC identity is controlled by cascades of temporal transcription factors. Although several cell-extrinsic cues regulate the timing of the neurogenesis-to-gliogenesis transition, few transcription factors involved in this process have been identified. A previous study suggested that Lhx2 regulates the neurogenesis-to-astrocytogenesis transition in the hippocampus but not in the cortex (Subramanian et al. 2011). The failure to identify Lhx2’s role in regulating the neurogenesis-to-gliogenesis transition in previous studies could result from: (i) Lhx2 functions was not inactivated at the appropriated timepoint to access gliogenesis onset. While conditional knockout of Lhx2 using Emx1-Cre (E10.5), Emx1-CreYL (E11.5), or Nestin-Cre (E11.5) leads to abnormal cortical morphology due to early corticogenesis defects, introducing Cre via IUE at E15.5 did not provide sufficient time for complete Lhx2 protein clearance before cortical gliogenesis onset (typically E16.5). (ii) Reliance on GFAP as a cortical gliogenesis marker. ALDH1L1, TNC, EGFR, and OLIG2—widely expressed in early glial cells—are detectable in E16.5 dorsal cortices, whereas GFAP is restricted to medial RGs and sparse astrocytes until postnatal stages (P1). However, given that Lhx2 plays a crucial role in promoting cortical neurogenesis (Mangale et al. 2008; Chou et al. 2009; Subramanian et al. 2011; Hsu et al. 2015), it is likely that Lhx2 also regulates the neurogenesis-to-gliogenesis transition in the cortex.

We conditionally knocked out *Lhx2* in cortical RGs after E13.5 using *hGFAP-Cre* (*hGFAP-Cre; Lhx2^F/F^,* or *Lhx2-*cKO) (Zhuo et al. 2001) without affecting its roles in cortical area patterning. Lhx2 inactivation increased GFAP in hippocampal/medial cortices but not dorsal-lateral cortices (Fig. 3A), consistent with prior findings (Subramanian et al. 2011). While ALDH1L1 and TNC were barely detectable in control E15.5 cortices, their expression was significantly upregulated in *Lhx2-cKO* cortices (Fig. 3A). Similarly, we observed an increase in EGFR and OLIG2 expression following *Lhx2* inactivation at E15.5 and E16.5 (Fig. 3A-B). The premature expression of late-stage RGs genes (ALDH1L1 and TNC) and the precocious generation of IGPs (EGFR+/OLIG2+ cells) indicated that *Lhx2* inactivation led to precocious gliogenesis. Consistent with this, neurogenesis was significantly reduced, as evidenced by decreased EOMES expression in E16.5 *Lhx2-* cKO mice (Fig. 3B). At P56, Lhx2-cKO mice exhibited a 28.3 ± 4.1% reduction in total cortical thickness relative to wild-type (WT) littermates (p = 0.002, unpaired t-test; n = 4 mice/genotype). While no significant changes were observed in FOXP1 (label Layer VI PyNs) and BCL11b (label Layer V PyNs) expression, CUX1 expression (label Layer II-IV PyNs) was seriously reduced in the *Lhx2-*cKO adult cortices (Fig. 3C), indicating that *Lhx2* inactivation after E13.5 selectively reduced the generation of upper-layer PyNs without affecting the production of deep-layer PyNs. In summary, the conditional knockout of *Lhx2* in cortical RGs during late neurogenic stages led to the premature generation of glia at the expense of upper layer PyNs.

**Fig. 3:**
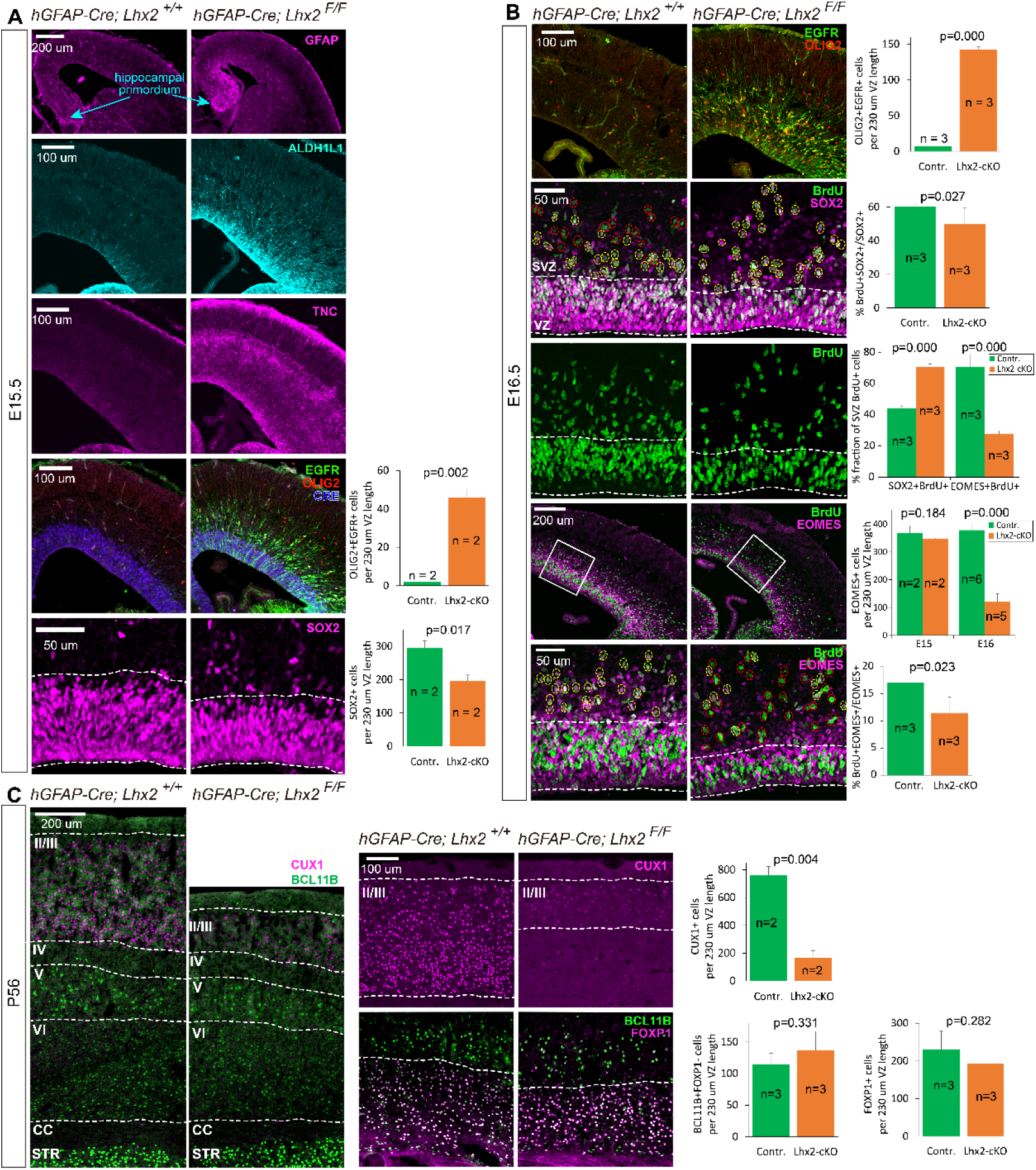
Lhx2 inactivation leads to premature gliogenesis at the expense of upper-layer PyNs. **A:** Immunostaining of GFAP, ALDH1L1, TNC, EGFR, OLIG2 and SOX2 in E15.5 control and *Lhx2* cKO cortices. **B:** Immunostaining of EGFR, OLIG2, SOX2, BrdU, and EOMES in E16.5 control and *Lhx2* cKO cortices. Yellow circles denote BrdU+ cells co-expressing SOX2 or EOMES; red circles indicate BrdU+ single-positive cells. **C:** Immunostaining of CUX1, BCL11B, and FOXP1 in P56 control and *Lhx2* cKO cortices. Quantification data represent mean ± SEM from n biologically independent animals. Statistical significance was assessed by unpaired Student’s t-test, with p-values indicated. **Abbreviations**: VZ, ventricular zone; SVZ, subventricular zone; II–VI, cortical layers II–VI; CC, corpus callosum; STR, striatum.

Upon *Lhx2* inactivation, scRNA-seq analysis of E15 FT/FACS-purified cortical progenitors from control and *Lhx2-*cKO mice confirmed a premature switch from neurogenesis to gliogenesis. A total of 9,758 cells were recovered from control cortices and 10,524 from *Lhx2-*cKO cortices. The cell types identified from the E15 control and *Lhx2-*cKO scRNA-seq data were comparable (Fig. S3A). Differentially expressed genes in each individual cell types between control and *Lhx2-*cKO mice were identified (Fig. S3D). Consistent with the increased ALDH1L1 and TNC protein levels, their mRNA expression in RGs also increased upon *Lhx2* inactivation (Fig. S3G). Moreover, the expression of many other late RG genes was upregulated upon *Lhx2* inactivation (Fig. S3E, H). GO terms that were downregulated over time in cortical RGs, such as those related to RNA processing, cell cycle, and chromatin remodeling (Fig. S2C), were also downregulated upon *Lhx2* inactivation (Fig. S3F). Thus, the RGs in E15 *Lhx2* mutants displayed molecular profiles characteristic of gliogenic-stage RGs.

Defects in stem cell maintenance were observed after *Lhx2* inactivation. The proportion of RGs decreased from 19.7% in control cortices to 11.4% in *Lhx2-*cKO cortices. The expression of *Pax6* and *Hes1*, both critical for stem cell maintenance, was reduced (Fig. S3G) (Sansom et al. 2009; Tuoc et al. 2009; Hatakeyama et al. 2004; Ochi et al. 2020). The GO term ’stem cell population maintenance’ was enriched in control RGs, while ’neurogenesis’ was enriched in *Lhx2-*cKO RGs (Fig. S3F, H). The reduction of RGs following *Lhx2* inactivation was supported by decreased SOX2+ cells (Fig. 3A-B). Thus, *Lhx2* inactivation resulted in a reduction of cortical RGs due to premature neuronal differentiation.

The percentage of cycling INPs (cINP-1 and cINP-2) decreased from 35.0% in the control cortices to 20.3% in the *Lhx2*-cKO cortices (Fig. S3B-C). Additionally, cell cycle gene expression levels were reduced in RGs upon *Lhx2* inactivation (Fig. S3F). The reduced proliferative capacity of both INPs and RGs was also supported by significantly fewer SOX2+ RGs and EOMES+ INPs incorporating BrdU following 30-minute pulse labeling in *Lhx2-cKO* versus *control* cortices (Fig. 3B). Thus, loss of *Lhx2* functions led to proliferation defects in both RGs and INPs.

In summary, LHX2 plays three primary roles during late neurogenesis: regulating the timing of the neurogenesis-to-gliogenesis transition, maintaining the stemness of cortical RGs, and promoting the proliferation of neuronal precursors.

### *Lhx2* Establishes an Active Epigenetic State to Promote Neurogenic Gene Expression

To elucidate how *Lhx2* regulates the cortical neurogenesis-to-gliogenesis transition, we performed LHX2 CUT&Tag-seq to identify its target genes. GO analysis of LHX2 target genes revealed that *Lhx2* regulates multiple aspects of cortical development, including neurogenesis, precursor proliferation, and cell fate commitment (Fig. 4A), consistent with the cortical defects observed upon *Lhx2* inactivation. In contrast, LHX2 exhibits minimal binding to late RG-specific or gliogenic genes, indicating that it does not directly regulate gliogenesis. Premature gliogenesis following *Lhx2* inactivation likely occurs secondarily due to the early cessation of neurogenesis. The majority of LHX2 binding sites were located within intronic (36.9%), downstream (19.2%), or intergenic (22.1%) regions, while only 17.1% were within promoter regions (Fig. S4A), suggesting that LHX2 preferentially binds to gene enhancers rather than promoters.

**Fig. 4:**
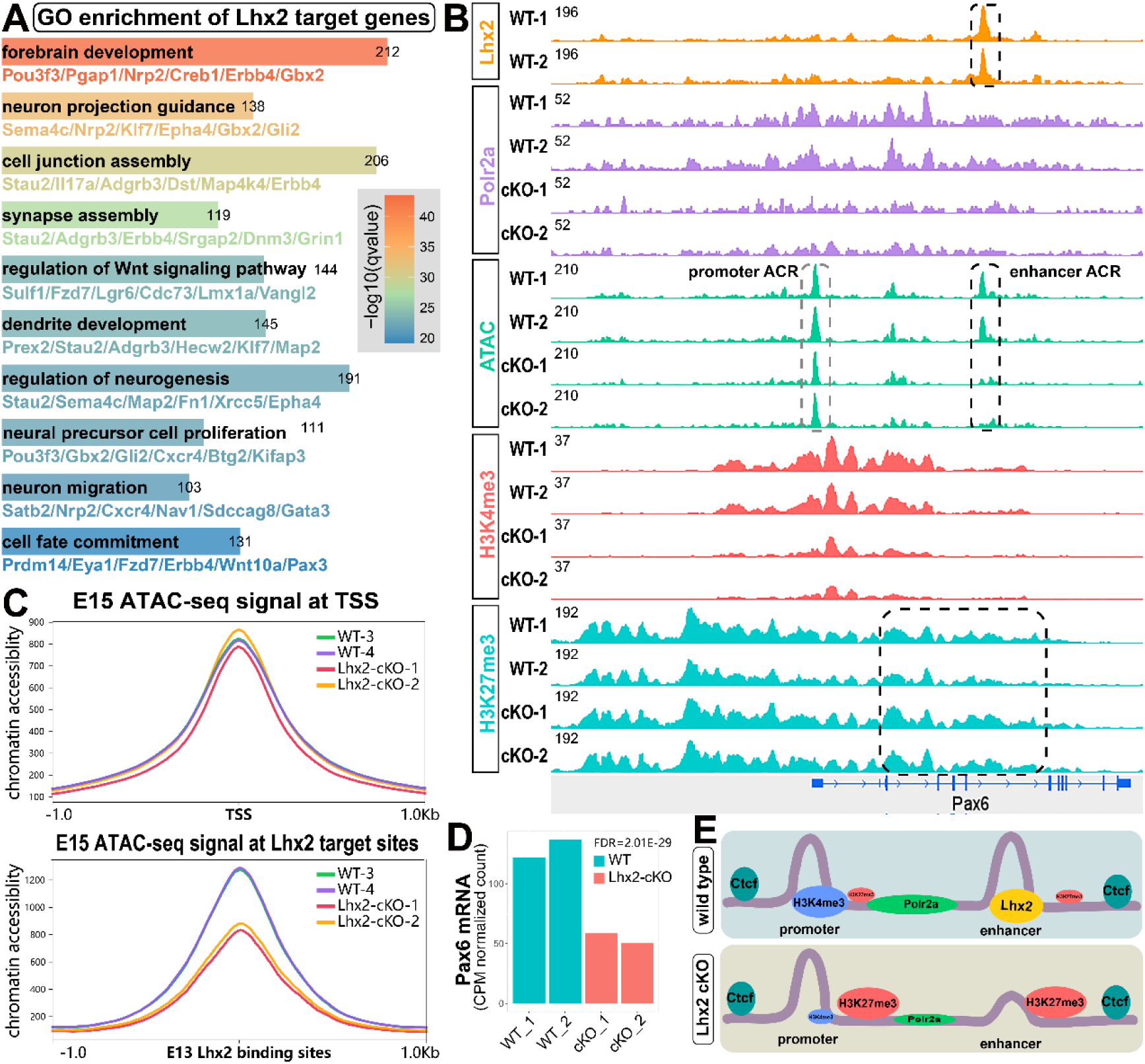
Lhx2 Inactivation Leads to Epigenetic Repression of Neurogenic Genes. **A**: GO enrichment analysis of Lhx2 target genes. **B**: Lhx2 inactivation led to epigenetic changes at the *Pax6* locus. Lhx2 CUT&Tag-seq from wild type E13 cortices; ATAC-seq, CUT&Tag-seq of Polr2a, H3K4me3, and H3K27me3, from E15 *control* and *Lhx2 cKO* cortices. **C**: Profile plots displaying ATAC-seq signal intensities surrounding transcription start sites (TSS) and LHX2 binding sites in both control and *Lhx2-cKO* cortical samples. **D:** Pax6 mRNA expression in both control and *Lhx2-cKO* cortical samples. FDR, False Discovery Rate. **E:** Schematic diagram illustrating the epigenetic repression induced by *Lhx2* inactivation.

To elucidate how LHX2 regulates its target gene expression, we profiled changes in the following epigenetic markers between E15 control and *Lhx2-cKO* cortices: (1) ATAC-seq to identify the accessible chromatin regions (ACRs); (2) CUT&Tag-seq for POLR2A, the largest subunit of RNA polymerase II, which is responsible for mRNA synthesis (Mita et al. 1995); and (3) CUT&Tag-seq for the active histone mark H3K4me3 and the repressive histone mark H3K27me3 Wiles and Selker, 2017). *Lhx2* inactivation resulted in reduced POLR2A occupancy at the *Pax6* locus (Fig. 4B), which is consistent with the decreased Pax6 mRNA expression (Fig. 4D). The Pax6 enhancer ACR, which overlap with the LHX2 binding site, significantly reduced accessibility upon *Lhx2* inactivation. In contrast, Pax6 promoter ACR, which not overlapping with LHX2 binding site, showed no change in chromatin accessibility (Fig. 4B). However, the H3K4me3 occupancy was greatly reduced at Pax6 promoters. While Pax6 locus is already highly occupied by H3K27me3, a slightly increase was observed upon *Lhx2-cKO*. Histone modifications can spread across large chromatin regions from initial transcription factor binding site, while the insulator *Ctcf* creates boundaries that insulate chromatin from epigenetic modifications (Narendra et al. 2015). CTCF binding sites segregate the Foxp4 locus into *Lhx2* related and *Lhx2* non-related chromatin regions (Fig. S4C). Similar changes in chromatin accessibility, POLR2A occupancy, and H3K4me3 and H3K27me3 modifications were observed in Lhx2-associated chromatin regions. Additionally, the Foxp1 locus exhibited a significant increase in H3K27me3 occupancy. In contrast, no changes were observed in Lhx2-independent chromatin regions.

We classified ACRs into *Lhx2*-target and *Lhx2*-nontarget ACRs by overlapping ATAC-seq peaks with LHX2 CUT&Tag-seq peaks and performed a genome-wide comparison of chromatin accessibility between *Lhx2-cKO* and control cortices. We recovered 51139 ACRs from the control and *Lhx2-*cKO cortical progenitors. Among these, 6905 were *Lhx2*-target ACRs (Fig. S4B). Upon *Lhx2* inactivation, 3,802 (55.1%) of these target ACRs exhibited reduced accessibility, 246 (3.6%) showed increased accessibility, and 2,857 (41.3%) remained unchanged (Fig. S4C). Notably, the ACRs with the most significantly reduced chromatin accessibility following *Lhx2-cKO* were consistently identified as *Lhx2*-target ACRs (Fig. S4C). While ATAC-seq signals surrounding transcription start sites (TSS) were unaffected, *Lhx2* inactivation induced a global decrease in chromatin accessibility specifically at *Lhx2*-target ACRs (Fig. 4C). These findings indicate that *Lhx2* inactivation leads to a widespread reduction in chromatin accessibility at its target ACRs.

In summary, *Lhx2* promotes the expression of neurogenic genes by establishing a transcriptionally permissive epigenetic state at its target loci (Fig. 4E). First, LHX2 binds to distal enhancers, enhancing local chromatin accessibility. Second, it regulates nearby promoters remotely by increasing H3K4me3 modification levels. Third, LHX2 suppresses H3K27me3 deposition across both local and adjacent chromatin regions, further facilitating gene activation.

### Temporal Dynamics DNA binding by LHX2 During Corticogenesis

*Lhx2* maintained neurogenic competence during late neurogenesis. Both *in situ* hybridization (ISH) and scRNA-seq revealed sustained *Lhx2* expression in cortical RGs throughout cortical development, with minimal temporal variation (Fig. 5A-B). This observation raises two mechanistic paradoxes: (1) Why do RGs progressively lose neurogenic competence despite persistent *Lhx2* expression, and (2) why does *Lhx2* fail to activate neurogenesis in gliogenic RGs, despite comparable expression levels in neurogenic and gliogenic populations? We propose that temporal shifts in LHX2’s DNA-binding dynamics drive the RG transition from neurogenesis to gliogenesis. To test whether Lhx2 exhibits stage-specific chromatin occupancy, we performed LHX2 CUT&Tag-seq on all E13 and E15 cortical cells, as well as on E18 FT/FACS-purified cortical progenitors. Consistent LHX2 CUT&Tag-seq signals were observed across all biological replicates (Fig. 5C). Both constitutively bound and developmentally dynamic LHX2 chromatin occupancy patterns were identified at the Pax6 and Cux1 loci (Fig. 5G). Strikingly, LHX2 exclusively interacted with a chromatin region downstream of *Pax6* at E13, whereas it progressively increased occupancy at the chromatin region within the second intron of Cux1 during E15-E18. We recovered 8435, 5980, and 5768 LHX2 binding sites (CUT&Tag-seq peaks) at E13, E15, and E18, respectively (Fig. 5D). The total number of LHX2 binding sites is progressively decreased over time, suggesting diminishing DNA-binding activity of Lhx2 during corticogenesis. Approximately 20% (a total of 2249) of those chromatin regions exhibited constitutive LHX2 occupancy across all three timepoints, while the majority interacted with LHX2 stage-specifically. Although LHX2 bound to distinct chromatin regions at different time points, its core DNA binding motif “TAATTA” remained unchanged (Fig. 5E-F). In summary, LHX2 exhibits temporally dynamic chromatin occupancy at distinct genomic loci during corticogenesis, accompanied by a progressive reduction in its overall DNA-binding activity.

**Fig. 5:**
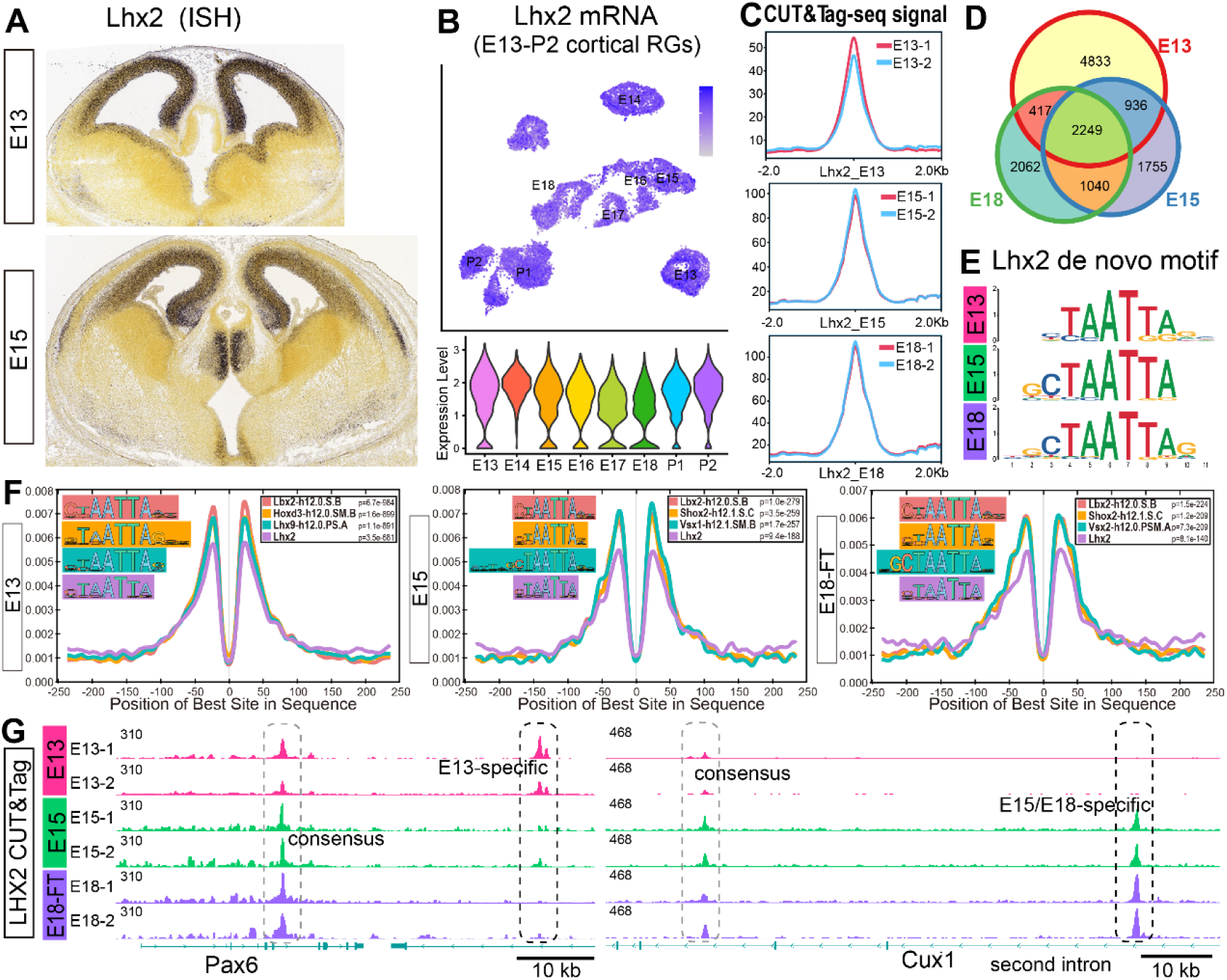
Lhx2 temporally binds to distinct genes to maintain chromatin accessibility. **A:** *In situ* hybridization of *Lhx2* in the embryonic brain (Allen Brain Atlas). **B**: *Lhx2* mRNA expression levels in cortical RGs across E13 to P2. **C**: Profile plots showing consistent LHX2 CUT&Tag-seq signal enrichment across all samples. **D**: Venn diagram showing the number of consensus and specific LHX2 binding sites across E13, E15, and E18 cortical progenitors. **E**: De novo motifs identified from E13, E15, and E18 LHX2 CUT&Tag-seq. **F:** Most significant motif in LHX2 chromatin occupancy regions. **G:** Track plots displaying consensus and dynamic Lhx2 occupancy at the Pax6 and Cux1 loci.

### Developmental Changes in Chromatin Accessibility Shape **LHX2** DNA Binding

The mechanisms underlying the temporally dynamic DNA binding of LHX2 remain unclear. This dynamic binding is unlikely to result from changes in LHX2 itself, as its binding motif remains unchanged during corticogenesis. Instead, it is more likely driven by changes in chromatin accessibility, as LHX2 tends to bind to accessible chromatin regions (ACRs) (Fig. S4B). To investigate chromatin accessibility changes between early and late RGs, we systematically analyzed single-nucleus ATAC-seq (snATAC-seq) data during cortical development. We performed snATAC-seq on E13 cortical cells and integrated the data with our previously published snATAC-seq data from E18 FACS-sorted hGFAP-GFP-expressing cells (Li et al. 2021) and publicly available snATAC-seq data from E14 cortices (Noack et al. 2022).

To determine cluster identities, we identified cell type-specific ACRs (Fig. S5A). Cluster annotations were then assigned based on these marker ACR profiles. The clusters identified in the snATAC-seq dataset were largely comparable to those in our scRNA-seq datasets, except for the absence of cINP clusters (Fig. 6A). While cell cycle genes exhibited cell type-specific expression, the ACRs surrounding them were detected similarly across all cell types, potentially explaining the lack of cINPs in the snATAC-seq dataset (Fig. S5A). As expected, the neurogenic gene *Neurog2* locus became progressively less accessible over time, whereas the gliogenic gene *Gfap* locus became more accessible in cortical RGs (Fig. 6B).

**Fig. 6:**
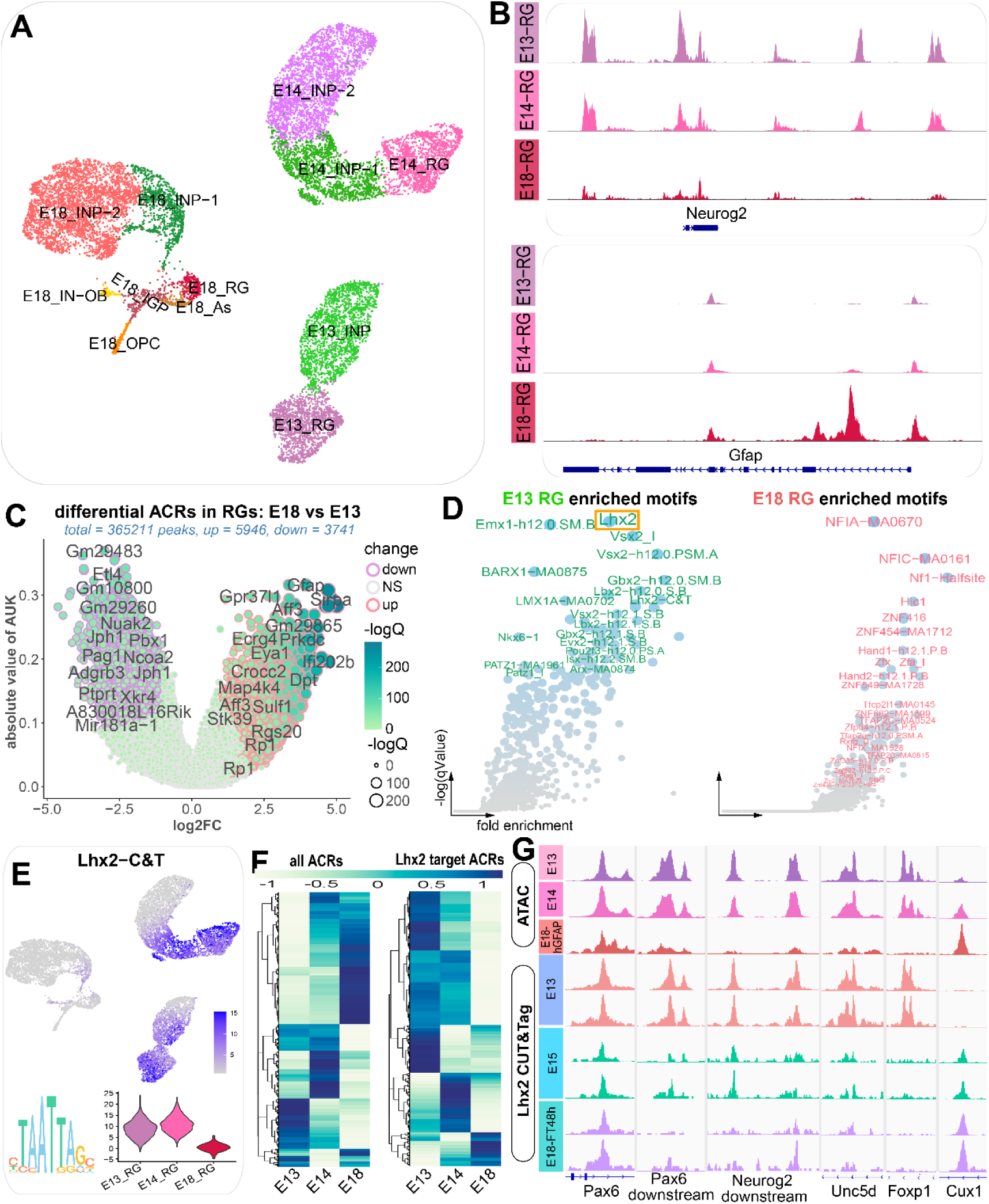
Time-Series snATAC-seq Analysis of Developing Cortex. **A**: UMAP projections of the combined snATAC-seq of cortical development (E13, E14, and E18). **B**: Track plots showing chromatin accessibility changes in RGs at the *Neurog2* and *Gfap* loci. **C**: Differential ATAC-peak analysis between E13 and E18 RGs. AUK, area under the Kappa curve; log2FC, log2(fold change); -logQ, -log10(q-value). **D**: Motifs enriched on E13 versus E18 RG specific ACRs. **E**: UMAP visualization of *Lhx2* motif activity in the developing cortex (upper) and violin plot displaying *Lhx2* motif activity in cortical RGs (lower). **F**: Heat maps showing the temporal changes in chromatin accessibility in cortical RGs. Notably, the majority of ACRs containing *Lhx2* binding motifs exhibit reduced accessibility over time (Right). **G**: Track plots displaying snATAC-seq peaks in RGs and Lhx2 CUT&Tag-seq peaks.

Each cell type opens specific chromatin regions, providing distinct binding sites for transcription factors. We computed transcription factor motif activities in each cell based on ACRs. As expected, motif activities for cell type-specific transcription factors, including Pax6, Olig2, Sox10, Sp9, and Neurog2, were detected in their respective cell types (Fig. S5B). To identify differentially enriched motifs between E13 and E18 RGs, we first identified E13- and E18-specific ACRs (Fig. 6C) and then compared the enriched motifs between these ACRs (Fig. 6D). Lhx2 motif activity was most enriched in E13 RG-specific ACRs (Fig. 6D). The three motifs recovered from our LHX2 CUT&Tag-seq data were merged into the Lhx2-C&T motif, confirming that Lhx2 motif activity was greatly reduced between early and late RGs (Fig. 6E). Consistent with reduced Lhx2 DNA binding, the majority of Lhx2 target ACRs became less accessible (Fig. 6F). At the Pax6 locus, the two Lhx2 target ACRs showed distinct accessibility patterns: while the consensus ACR retained Lhx2 occupancy, the downstream ACR became inaccessible in late RGs and lost Lhx2 binding. Similar changes were observed at the neurogenic gene Neurog2, the PyN migration gene Unc5d, and the layer VI PyN-specific gene Foxp1. In contrast, the layer II PyN gene Cux1 exhibited progressively increased chromatin accessibility, accompanied by enhanced Lhx2 occupancy (Fig. 6G).

In summary, changes in Lhx2 chromatin occupancy closely aligned with changes in chromatin accessibility, indicating that dynamic Lhx2 DNA binding results from developmental changes in chromatin accessibility.

### IGPs Sequentially Open Distinct Chromatin Regions for Binding Cell Fate-Determining Transcription Factors

Our snATAC-seq data revealed that distinct cell types open specific chromatin regions, enabling the binding of transcription factors associated with cell fate determination. To identify putative transcription factors that specify each cell fate in the glial lineage, we analyzed transcription factor motif activities along the glial lineage development.

The glial lineage trajectories reconstructed from the E18 *hGFAP-GFP* snATAC-seq data revealed that IGPs first produce astrocytes, followed by oligodendrocytes and olfactory bulb (OB) interneurons, consistent with the glial lineage trajectories inferred from the scRNA-seq data (Fig. 7A). Motif activities for genes such as *Sox10* and *Dlx5* were specifically detected in oligodendrocytes and OB interneurons, consistent with their expression patterns (Fig. 7H, J). Interestingly, we observed many genes that are broadly expressed but exhibit lineage-specific motif activity. For example, while the mRNAs of *Nfia, Nfib,* and *Rbpj* are detected in all glial lineage cells, their motif activities are restricted to the astrocyte lineage (Fig. 7B-D). Motif activities of *Nfia* and *Nfib* were specifically detected in young IGPs around the astrocyte bifurcation points, suggesting their roles as astrocyte fate determinants. In contrast, *Rbpj* motif activity was primarily detected in RGs and astrocytes but not in young IGPs at the astrocyte bifurcation points, indicating its role in maintaining RG and astrocyte identity rather than determining astrocyte fate. *Sox2* motif activity was detected in both young and old IGPs, while *Ascl1, Olig2*, and *Tcf4* motif activities were observed in old IGPs after the production of astrocytes (Fig. 7E-G). While *Olig2* motif activity was higher in cells progressing toward oligodendrocytes, *Tcf4* motif activity was higher in cells progressing toward OB interneurons. Additionally, Smad2 and Meis2 motif activities were specific higher in oligodendrocytes and OB interneurons, respectively (Fig. 7I, K). Motifs containing the core sequences CAGCTG|CACCTG|CAGGTG|ACAATG, GGAA|TCA-TCA, and GATA|TTGT were enriched in IGPs, astrocytes, and oligodendrocytes, respectively. Furthermore, AT-rich motifs were specifically enriched in OB interneurons (Fig. 7L).

**Fig. 7:**
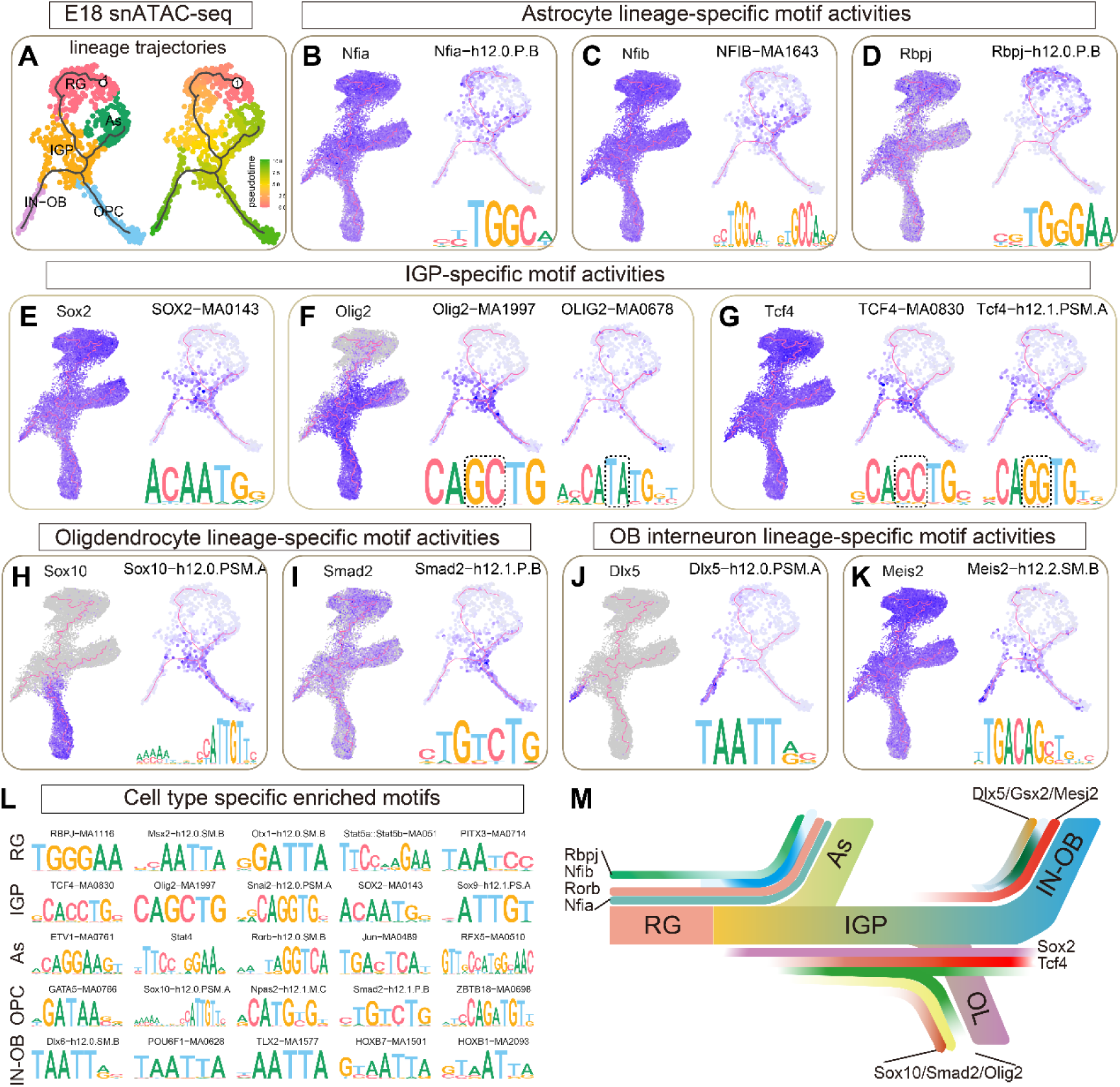
Dynamic transcription factor motif activities along glial lineage cells. **A:** UMAP visualization of E18 snATAC-seq glial lineage cells with lineage trajectories, colored by cell type (left) and pseudotime (right). **B-K:** A comparison of gene expression pattern with its corresponding motif activity in glial lineage cells. In each panel, **left**: mRNA expression levels of representative genes in glial lineage cells; **right**: motif activities of the corresponding transcription factors in glial lineage cells. **L:** Top overrepresented motifs for each cell types. **M:** Schematic diagram illustrating transcription factor activities along the glial lineage.

Collectively, these findings indicate that young IGPs open chromatin regions to allow the binding of *Nfia* and *Nfib*, thereby specifying astrocyte fate. In contrast, old IGPs open chromatin regions for the binding of *Olig2* and *Tcf4*, facilitating the generation of oligodendrocytes and OB interneurons (Fig. 7M).

## Discussion

During cortical development, RGs transition from early to late stages, sequentially producing INPs and IGPs. While INPs exclusively differentiate into PyNs, IGPs transition from young to old, sequentially generating cortical astrocytes, oligodendrocytes, and OB interneurons. As RGs transition from early to late and IGPs progress from young to old, they sequentially express distinct genes to coordinate production of distinct cell types. Our analysis also uncovered that chromatin accessibility plays a critical role in regulating the temporal patterning of RGs and IGPs by restricting the functions of broadly expressed transcription factors to specific periods and particular cell types. We identified Lhx2 as a temporal transcription factor that epigenetically regulates the timing of the neurogenesis-to-gliogenesis transition.

### Cell Intrinsic Programs Regulates the Neurogenesis-to-Gliogenesis Transition

Epigenetic changes in RGs significantly contribute to cell-intrinsic programs that govern the developmental switch in cellular competence in cortical RGs. Previous studies suggest that neurogenic genes are gradually inactivated by increasing trimethylation at H3K27 (Hirabayashi et al. 2009), while gliogenic genes are gradually activated by DNA demethylation (Takizawa et al. 2001b; Fan et al. 2005b; Sanosaka et al. 2017). Our findings show that temporal changes in chromatin accessibility play a key role in regulating RG competence transition. *Lhx2* plays a crucial role in promoting cortical neurogenesis (Mangale et al. 2008; Chou et al. 2009; Subramanian et al. 2011; Hsu et al. 2015). However, RGs transition from neurogenesis to gliogenesis, despite constant Lhx2 expression. As corticogenesis proceeds, the accessibility of neurogenic genes decreases, while that of gliogenic genes increases. Developmental changes in chromatin accessibility prevent LHX2 from binding to neurogenic genes in late-stage RGs, thereby inhibiting LHX2 induced neurogenesis. To summarize, cell-intrinsic changes in DNA methylation, histone modification, and chromatin accessibility contribute to the neurogenesis-to-gliogenesis transition. It is likely that the epigenetic states of RGs endow them with either neurogenic or gliogenic competence.

### Bivalent epigenetic regulation modulates chromatin accessibility of *Lhx2* target ACRs

We hypothesize that *Lhx2* target ACRs are subject to a dynamic balance between active and repressive epigenetic factors for the following reasons. First, the majority of E13 *Lhx2* target ACRs become closed in late-stage cortical RGs (e.g., the *Unc5d* locus in Fig. 7a), indicating that negative epigenetic regulation gradually surpasses the positive regulation by *Lhx2* at those ACRs. Second, *Lhx2* interacts directly or indirectly with both the NuRD and BAF complexes. Mass spectrometric analysis of *Lhx2*-interacting proteins revealed co-immunoprecipitation of NuRD subunits (Rbbp4 and Ss18l1) and BAF subunits (Smarcc1, Smarcc2, and Smarce1) with Lhx2 (Muralidharan et al. 2017; Nguyen et al. 2022). Both NuRD and BAF complexes are ATP-dependent chromatin remodeling complexes. NuRD is a transcriptional corepressor that couples ATP-dependent chromatin remodeling with histone deacetylase activity (Nitarska et al. 2016; Bornelöv et al. 2018), while the BAF complex primarily activates gene expression by evicting nucleosomes from ACRs (Brahma and Henikoff 2024; Narayanan et al. 2015). Third, *Lhx2* and Polycomb complexes exert opposing effects on the chromatin accessibility of neurogenic genes (Hirabayashi et al. 2009). *Lhx2* positively regulates the chromatin accessibility of neurogenic genes to maintain neurogenic competency for producing upper layer PyNs, while Polycomb complexes negatively regulate these genes to facilitate the transition to gliogenesis. Thus, a balance between positive and negative epigenetic regulation delicately controls the chromatin accessibility of *Lhx2* target ACRs.

Epigenetic factors are usually recruited to specific chromatin regions via interactions with transcription factors. At *Lhx2* target ACRs, active epigenetic factors can be recruited by *Lhx2*, which recognizes specific DNA motif. Repressive epigenetic factors are recruited to specific genes by yet unidentified factors. It is unlikely that *Lhx2* recruits both active and repressive epigenetic factors, as *Lhx2* primarily positively regulates its target ACRs. Alternatively, a putative *Lhx2* "competitor" may be required for the accurate targeting of repressive epigenetic factors, which remains to be identified in the future.

### Dynamic chromatin accessibility restricts the functions of broadly expressed transcription factors to specific developmental periods and cell types

Chromatin DNA is tightly packaged into higher-order structures, largely occupied by nucleosomes, leaving only a limited number of regions accessible for physical interactions with transcription factors and other chromatin-associated proteins (Klemm et al. 2019). These ACRs can function as enhancers, promoters, and insulators, cooperatively regulating gene expression. Transcription factors, essential for gene regulation, modulate transcriptional activity by binding to specific DNA sequences (Spitz and Furlong 2012). Nucleosomes and higher-order chromatin packaging form a physical barrier to transcription factor binding due to the tight wrapping of DNA around histone proteins. Thus, most transcription factors bind preferentially to ACRs, with the exception of a few pioneer factors. Changes in chromatin accessibility can alter the physical interactions between chromatin DNA and transcription factors, enabling a single transcription factor to function in a context-dependent manner.

*Lhx2* exerts multiple functions at distinct developmental stages during corticogenesis. At the onset of neurogenesis, *Lhx2* promotes cortical primordium identity by repressing the hem and anti-hem fate (Chou and Tole 2019). During early neurogenesis, *Lhx2* promotes the specification of layer 6 over layer 5 PyN identity (Muralidharan et al. 2017). During late neurogenesis, *Lhx2* maintains neurogenic competence for generating upper-layer PyNs and prevents premature gliogenesis. Consistent with its dynamic roles, *Lhx2* shows stage-specific DNA-binding activity during corticogenesis. In cortical RGs, *Lhx2* binds to distinct sets of genes at different developmental stages, with its overall DNA-binding activity progressively decreasing over time. The dynamic DNA-binding activities of *Lhx2* coordinate the switch from neurogenic to gliogenic potential in cortical RGs. Initially, *Lhx2* binds to neurogenic genes, maintaining neurogenic competence in cortical RGs. As corticogenesis progresses, Lhx2 DNA-binding activity gradually decreases, leading RGs to lose neurogenic competence and transition to gliogenesis.

In *Drosophila*, the temporal identity of NSCs is endowed by cascades of temporal transcription factors whose expression varies over time. Our study highlights the crucial role of chromatin accessibility dynamics in temporal patterning. RGs temporally open distinct chromatin regions, enabling stage-specific DNA binding by transcription factors that are expressed consistently without temporal variation. This allows the cells to respond differently to the same factors at different developmental stages. Despite the temporal patterning of RGs, chromatin accessibility dynamics may also contribute to the diversification of IGPs into astrocytes, oligodendrocytes, and OB interneurons. These findings raise important questions for future exploration: How is stage- or cell type-specific chromatin accessibility established? In particular, how do neurogenic genes progressively lose chromatin accessibility, while gliogenic genes gain it over time?

## Methods

### Mice

All experiments performed at Fudan University Shanghai Medical College were approved by Fudan University Animal Ethics Committee. *Lhx2*-flox mice (Strain NO. T036697) were purchased from GemPharmatech (Nanjing, China). *hGFAP-Cre* mice and *hGFAP-GFP* mice were previously described (Zhuo et al., 2001). The transgenic mice were genotyped by PCR using the following primers: *Lhx2*, “TGGTCTAAGGATCAAGAGGCTAC”, “GCGGTTAAGTATTGGGACAGAG”; *hGFAP-Cre*, “ACGAGTGATGAGGTTCGCAAGA”, “GATTAACATTCTCCCACCGTCAGT”. The day of vaginal plug detection was designated E0.5. The day of birth was designated P0.

### Immunohistochemistry and imaging

Adult mice were perfused with 1× PBS (pH 7.4), followed by 10min 4% paraformaldehyde (PFA) in PBS. The adult brain tissues were postfixed overnight in 4% PFA. Embryonic brains were immersion fixed for 4 hours in 4% PFA. To assess progenitor proliferation capacity, pregnant dams carrying E16.5 embryos received intraperitoneal injections of 5-bromo-2′-deoxyuridine (BrdU; 50 mg/kg in 0.9% NaCl) 30 minutes prior to sacrifice. All the brain tissues were cryoprotected overnight in 30% sucrose, frozen in the embedding medium, and cryosectioned at 14/30 μm using a cryostat (CM1950, Leica, USA). The sections were first permeabilized with 0.05% Triton X-100 for 30 min, followed by incubation in blocking buffer (5% donkey serum and 0.05% Triton X-100 in TBS) for 2 h. The sections were incubated with primary antibodies overnight at 4°C. Primary antibodies were removed by washing three times with TBS. The sections were incubated with secondary antibodies for 2 hours at room temperature. The secondary antibodies were from Jackson ImmunoResearch and Invitrogen. Finally, the sections were counterstained with DAPI for 3 min before being mounted in fluorescence mounting medium (DAKO S3023). The following primary antibodies were used: OLIG2 (1:1,000, rabbit, Millipore AB9610), OLIG2 (1:1,000, antigen retrieval, mouse, Millipore MABN50), EGFR (1:2,000, goat, R&D System AB_355937), ALDH1L1 (1:1,000, rabbit, antigen retrieval, Abcam ab10712968), BCL11B (1:1500, rat, Abcam, ab18465), EOMES (1:300, rabbit, Abcam, ab23345), EOMES (rat, 1:500, Thermo Fisher, 12-4875-82), FOXP2 (1:500, goat, Santa Cruz, sc-21 069), CUX1 (1:200, rabbit, Santa Cruz, sc-13024), TNC (1:200, rabbit, Abcam, ab108930), BrdU (1:200, Rat, Accurate Chemical, OBT0030s).

The fluorescent images were taken with Olympus FV1000 confocal microscope system using a 10×,20×, or a 40× objective. Z-stack confocal images were reconstructed using the FV10-ASW software. All images were merged, cropped, and optimized in Photoshop CC without distortion of the original information.

### Cortical tissues dissociation

The embryonic cortices were dissected in ice-cold PBS and digested for 15-20 mins at 37◦C with 80 U/mL papain (sigma, p3125), 200 U/mL DNase I (Invitrogen,18047019) in DMEM cell culture medium (Gibco, 10566024). The tissues were dissociated into single cell suspension by pipetting several times. FBS containing DMEM was added to stop the enzymatic reaction. The cells were filtered with a pre-wet 40 μM cell strainer (Corning, 352340) and pelleted at 300 g (RCF) for 5 min. The cells were resuspended in 10% FBS containing DMEM. Viable cells were counted using 0.4% Trypan Blue under microscope.

### Bulk ATAC-seq and bulk RNA-seq

RNA-seq libraries were constructed from 20,000 to 50,000 dissociated cortical cells using the Hieff NGS® Ultima Dual-mode RNA Library Prep Kit (YEASEN, Cat. #12309) following the manufacturer’s instructions.

For bulk ATAC-seq, 20,000 to 50,000 cells were pelleted at 500 × g for 3 minutes. We added 20 μL of the Tn5 tagmentation mixture to simultaneously facilitate cell/nuclear membrane penetration and Tn5 tagmentation. The Tn5 tagmentation mixture consisted of 6.6 μL H2O, 7 μL PBS, 0.2 μL 10% Tween-20, 0.2 μL 1% digitonin, 4 μL 5× Tn5 buffer, and 2 μL of ME adapter sequence-loaded Tn5 transposases (Vazyme, TD501). The DNA fragments were purified using 2X VAHTS DNA Clean Beads (Vazyme, N411-01). PCR (12–15 cycles) was used to amplify the Tn5-fragmented DNA and incorporate i5/i7 adapters, sequencing primers, and sample indices for Illumina sequencing. The 200–800 bp PCR products were purified using 0.6×/1.2× SPRIselect Reagent (Beckman Coulter, B23318) and subsequently subjected to Illumina PE150 Nova sequencing.

### CUT&Tag-seq

For CUT&Tag-seq, 1–2 × 10⁵ cells were washed with 500 μL of wash buffer and centrifuged at 600 g for 3 minutes at room temperature. The cell pellets were then resuspended in 100 μL of wash buffer. Concanavalin A-coated magnetic beads were washed twice with 100 μL of binding buffer. Next, 10 μL of the activated beads were added to the cell suspension and incubated at room temperature for 10 minutes. Following incubation, the binding buffer was removed, and the bead-bound cells were resuspended in 50 μL of antibody buffer. Subsequently, 1 μg of primary antibodies was added and the mixture was incubated overnight at 4°C. The following primary antibodies were used: Lhx2 (Abcam, ab184337), H3K4me3 (ABclonal, A22146), H3K27me3 (ABclonal, A22396), and Polr2a (BioLegend, 664906). After the removal of the primary antibody solution, 0.5 μg of the appropriate secondary antibodies were added to 50 μL of Dig-wash buffer and incubated at room temperature for 1 hour. The cells were then washed three times with Dig-wash buffer to remove any unbound antibodies. Next, the cells were incubated with 0.04 μM Tn5 transposase in 100 μL of Dig-300 buffer at room temperature for 1 hour. The cells were washed three times with Dig-300 buffer to remove any unbound transposase. The cells were then resuspended in 300 μL of tagmentation buffer and incubated at 37°C for 1 hour. To terminate tagmentation, 10 μL of 0.5 M EDTA, 3 μL of 10% SDS, and 2.5 μL of 20 mg/mL Proteinase K were added to the 300 μL sample, followed by overnight incubation at 37°C. DNA was purified using phenol–chloroform–isoamyl alcohol extraction, followed by ethanol precipitation and RNase A treatment. PCR amplification and Illumina sequencing were performed using a protocol similar to that of ATAC-seq.

### Data analysis of bulk RNA-seq, bulk ATAC-seq and CUT&Tag-seq

Raw FastQ files were processed using Trim Galore to remove low-quality bases and adapter sequences. Cleaned reads were then aligned to the mouse reference genome GRCm39 (GENCODE release M31) using Subread. For RNA-seq analysis, gene-level read counts were quantified using featureCounts, followed by differential expression analysis with edgeR. For ATAC-seq and CUT&Tag-seq data, we removed PCR duplicates, unpaired reads, and low-quality alignments (MAPQ < 30) from BAM files using Sambamba and SAMtools. The cleaned BAM files were then converted into RPKM-normalized bigWig files using deepTools. The bigWig files were visualized in IGV (v2.16.2) with “Group Autoscale” to enable comparative analysis across samples. We performed peak calling using two independent methods: MACS3 (v3.0.1) for analyzing BAM files from all biological replicates (n=2), and LanceOtron (v1.2.6) for analyzing merged bigWig files. The peaks were annotated to the nearest genes using ChIPseeker.

For differential ATAC-peak analysis between control and *Lhx2* cKO cortical progenitors, peaks from both control and mutant ATAC-seq data were merged into common peaks using the reduce function in GenomicRanges. Read counts within these common peaks were obtained from the cleaned BAM files using featureCounts in the Rsubread package. Differential analysis of read counts was conducted using EdgeR.

The consensus and temporal dynamics of *Lhx2* binding sites were identified using Vennerable and VennStem. Overlaps between *Lhx2* CUT&Tag-seq peaks and snATAC-seq peaks were identified using the findOverlaps function in GenomicRanges, while overlaps between *Lhx2* CUT&Tag-seq peaks and bulk ATAC-seq peaks were determined using BEDTools intersect.

### scRNA-seq and snATAC-seq

For scRNA-seq, 1–2 × 10⁴ cells were loaded into the 10X Genomics Chromium platform to recover between 6,000 and 12,000 cells. scRNA-seq libraries were constructed using the Chromium Next GEM Single Cell 3’ Reagent Kits v3.1, following the manufacturer’s instructions (manual document part number: CG000205 Rev C, 10x Genomics). The cDNA libraries were then purified, quantified using an Agilent 2100 Bioanalyzer, and sequenced on an Illumina NovaSeq 6000 system.

For snATAC-seq, 2–5 × 10⁵ cells were subjected to nuclei isolation according to the manufacturer’s instructions (manual document part number: CG000366 Rev D, 10x Genomics). A total of 1–2 × 10⁴ transposed nuclei were loaded to recover 3,000– 6,000 cells. snATAC-seq libraries were constructed using the Chromium Next GEM Single Cell ATAC Reagent Kits v1.1, following the manufacturer’s instructions (manual document part number: CG000209 Rev F, 10x Genomics). The snATAC-seq libraries were sequenced on an Illumina NovaSeq 6000 system, yielding approximately 13,000–30,000 read pairs per cell.

### Data Analysis of scRNA-seq

To generate the gene expression matrix for scRNA-seq, we used Cell Ranger count (10x Genomics Cell Ranger v7.1.0) with the mouse genome GRCm39 as the reference. The expression matrices were corrected for ambient mRNA contamination using SoupX. Low-quality cells were filtered out based on total counts, the number of detected genes, and the percentage of mitochondrial genes. Doublets were identified and removed using DoubletFinder. Cells not belonging to cortical radial glial (RG) lineages were also excluded. The cleaned count matrices were merged into two datasets corresponding to neurogenesis and gliogenesis. Seurat objects were created from these datasets (https://satijalab.org/seurat/articles/install.html). The data were log-normalized and scaled, with cell cycle genes and mitochondrial genes regressed out. Principal component analysis (PCA) was performed, and batch effects were corrected using Harmony integration. UMAP dimensionality reduction and clustering were then performed on the Harmony-integrated PCs. Differential expression analysis between neurogenic and gliogenic RGs was conducted using both EdgeR and presto.

The count matrix specific to cortical RGs was subset from the scRNA-seq datasets. Counts were normalized by sequencing depth for each cell. Genes with low expression across all time points (max mean < 0.3) were filtered out. To generate the age-related gene expression matrix, we computed the average gene expression at each time point, ordering them by age. Genes with minimal variation (standard deviation > 0.2) were also filtered out. Gene temporal dynamic patterns were clustered using Mfuzz. Gene Ontology (GO) enrichment and Gene Set Enrichment Analysis (GSEA) were performed using ClusterProfiler, with similar GSEA GO terms grouped using aPEAR.

The Seurat object was converted into Monocle object to reconstruct lineage trajectories and infer cell pseudotime (https://cole-trapnell-lab.github.io/monocle3/). Genes differentially expressed along the pyramidal neuron (PyN) or glial lineage trajectories were identified using the Moran’s I test. Expression levels for each gene were fitted with the lineage pseudotime using the generalized additive model (GAM) in the mgcv package, and fitted values were extracted. Pseudotime-dynamic genes were identified using Mfuzz and clustered into co-expression modules.

Unspliced and spliced reads were annotated using Velocyto, which distinguishes between nascent pre-mRNA and mature mRNA in scRNA-seq datasets. Cellular dynamics were inferred from mRNA splicing kinetics using scVelo. Cell state transitions were reconstructed by integrating RNA velocity with Monocle3 pseudotime analysis using CellRank2.

### Data Analysis of snATAC-seq

The Cell Ranger ATAC pipeline (v2.1.0) was used to process the FastQ files from snATAC-seq. To combine snATAC-seq datasets across all time points, a common set of distinct, non-overlapping peaks was created by merging all intersecting peaks using the reduce function in GenomicRanges. Counts within these common peaks were quantified for each cell to create Signac objects for each sample (https://stuartlab.org/signac). The Signac objects were then merged and annotated with mouse genome annotation GRCm39-M31. Cells were filtered based on total counts, nucleosome signal, TSS enrichment, and peak enrichment. The Signac object underwent TF-IDF normalization, Latent Semantic Indexing (LSI) dimensional reduction, UMAP reduction, and clustering. We identified cluster-specific peaks using FindAllMarkers and annotated cell clusters based on the characteristic peaks of their marker genes. Differential peak analysis between E13 and E18 RGs was performed using both EdgeR and presto.

Motif activities per cell were calculated using motifs from mouse transcription factors and chromatin regulators, which were downloaded from JASPAR, CIS-BP, Homer, and HOCOMOCO (Rauluseviciute et al. 2024; Weirauch et al. 2014; Vorontsov et al. 2024). After filtering motifs with low information content (IC) scores and merging similar motifs associated with the same genes, a total of 2,858 motifs related to 1,200 genes were retained. Per-cell motif activity scores were computed using chromVAR, and cell type-specific motif activities were identified.

## Acknowledgments

This study was supported by the Ministry of Science and Technology of China (STI2030-2021ZD0202300), National Natural Science Foundation of China (NSFC 31820103006, 32070971, 32100768, 32200776, and 32200792), Shanghai Municipal Science and Technology Major Project (No. 2018SHZDZX01), ZJ Lab, and Shanghai Center for Brain Science and Brain-Inspired Technology.

## Author contributions

X.L. designed research. L.Y. performed research. Y.G., Z.W., Z.L., G.L., Z.X., Y.Y., Z.Y., and X.L. contributed new reagents/analytic tools; X.L. performed data analysis and wrote the paper.

## Data availability

scRNA-seq, snATAC-seq, Bulk ATAC-seq, Cut&Tag-seq data used in this study have been deposited in the Gene Expression Omnibus (GEO) and are accessible through GEO Series accession number GSE298893. The E14 cortical scRNA-seq was from previous studies (GSE155677) (Noack et al. 2022). scRNA-seq data of E18 FT/FACS cortical progenitors (GSE221389) and snATAC-seq data of E18 *hGFAP-GFP* cortical progenitors (GSE161132) was used in our previous studies (Li et al. 2021, 2023). The code used for generating the data and all the figures are available under https://github.com/xiao00su/temporal.

