## supplemental figures Fig. S1-S5 for "Mouse Cortical Cellular Diversification Through Lineage Progression of Radial Glia"

**Supplementary Figure Legends**

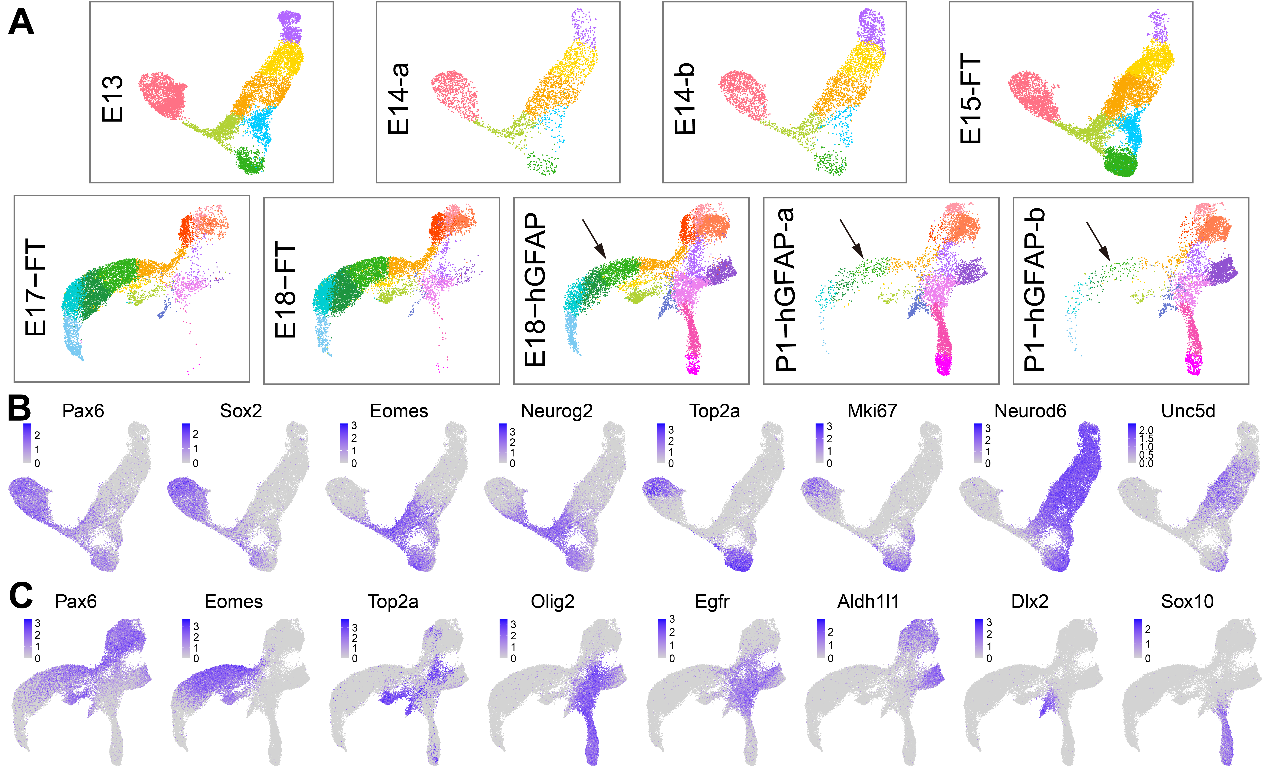
Fig. S1:  **Cell type marker expression in the neurogenic and gliogenic datasets**. **A**: UMAP visualization of each sample, with arrows indicating a sharp reduction in PyN lineage cells between E18 and P1. **B-C**: Feature plots of representative cell type markers of neurogenic (B) and gliogenic (C) datasets.

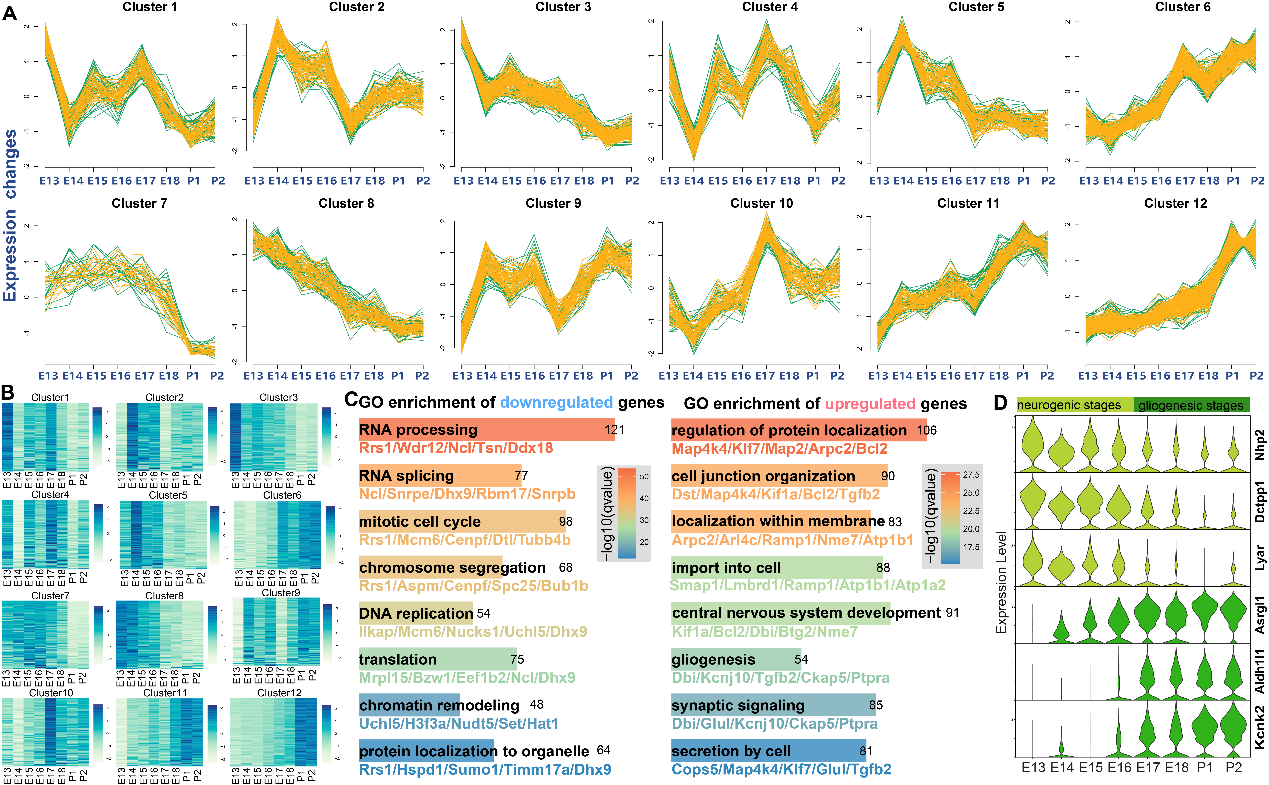

Fig. S2: **Temporal Dynamics of Gene Expression in Cortical RGs**. **A**: Clustering of the dynamic patterns of temporal gene expression levels in cortical RGs. **B**: Heatmap plotting of gene expression levels for each gene cluster, colored by average relative counts. **C**: GO enrichment analysis of temporally downregulated (left) and upregulated (right) genes. **D**: Violin plots showing genes primarily expressed in RGs during neurogenic (light green) and gliogenic (dark green) stages.

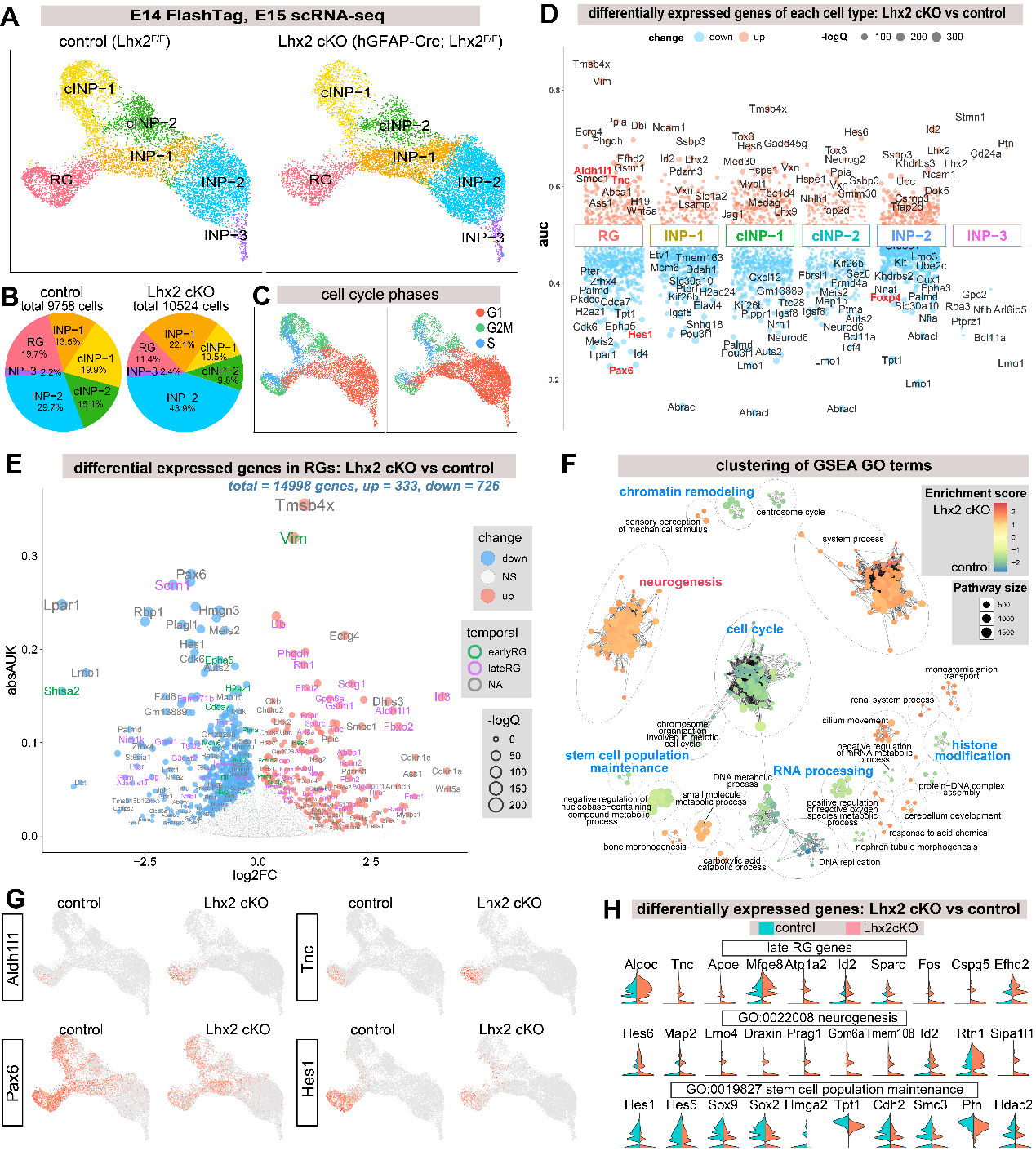
Fig. S3: **scRNA-seq analysis of FT/FACS-sorted cortical progenitors from control and Lhx2 cKO mice. A**: UMAP projections of the combined scRNA-seq data from control or Lhx2 cKO cortical progenitors, colored by cell types. **B**: Comparison of cell type composition between control and Lhx2 cKO mice. **C**: UMAP plot showing cell cycle phase identities. **D**: Differential gene expression analysis for each cell type between control and Lhx2 cKO mice. **E**: Differential gene expression between control and *Lhx2*-cKO cortical RGs. Many late-stage RG genes (purple) are predominantly upregulated in cortical RGs upon *Lhx2* inactivation. **F**: Clustering of GSEA GO terms that enriched in control and Lhx2-cKO RGs, colored by enrichment scores. Lower enrichment scores indicate GO term enrichment in control RGs, while higher scores indicate enrichment in *Lhx2*-cKO RGs. Each dot representing a GO term and similar GO terms are grouped into clusters. **G**: Representative differentially expressed genes between control and Lhx2 cKO mice. **H**: Split violin plots illustrating differential gene expression between control and *Lhx2* cKO RGs.

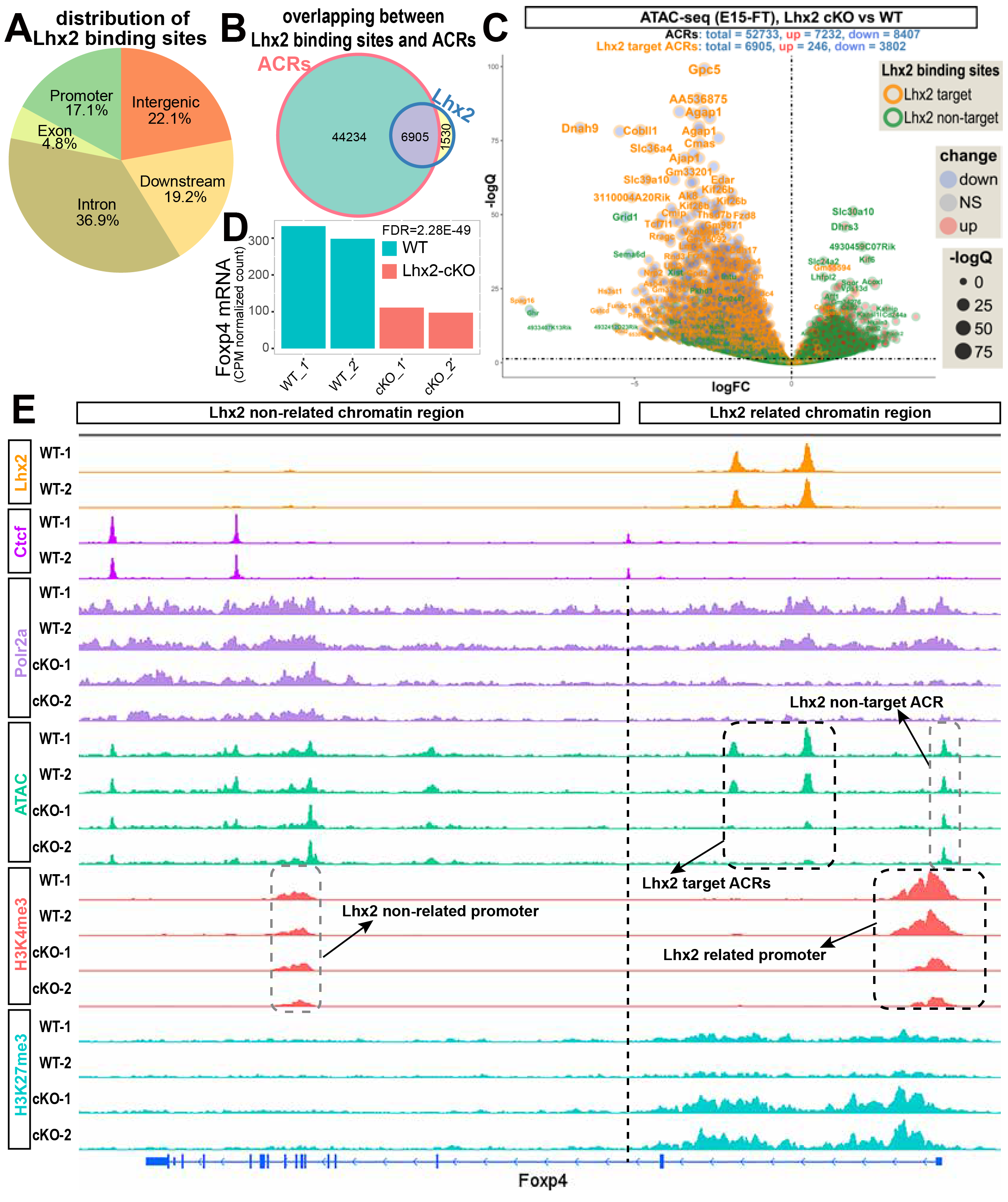
Fig. S4: **Lhx2 increases local chromatin accessibility but remotely regulates gene promoters within its associated chromatin regions**. **A**: A pie chart showing fractional distribution of Lhx2 binding sites. **B**: A Venn diagram showing the overlap between Lhx2 binding sites and ACRs (left), and a pie chart showing fractional distribution of Lhx2 binding sites (right). **C**: Differential ATAC-peak analysis between E15 control and Lhx2 cKO cortical progenitors, colored by Lhx2 target/non-target ACRs. **D**: Foxp4 mRNA expression in both control and *Lhx2-cKO* cortical samples. **E:** Enrichment of H3K4me3, H3K27me3, and Polr2a at the *Foxp4* locus in *control* and *Lhx2 cKO* cortical cells. The dashed line indicates the boundary between Lhx2-related and non-related chromatin regions.

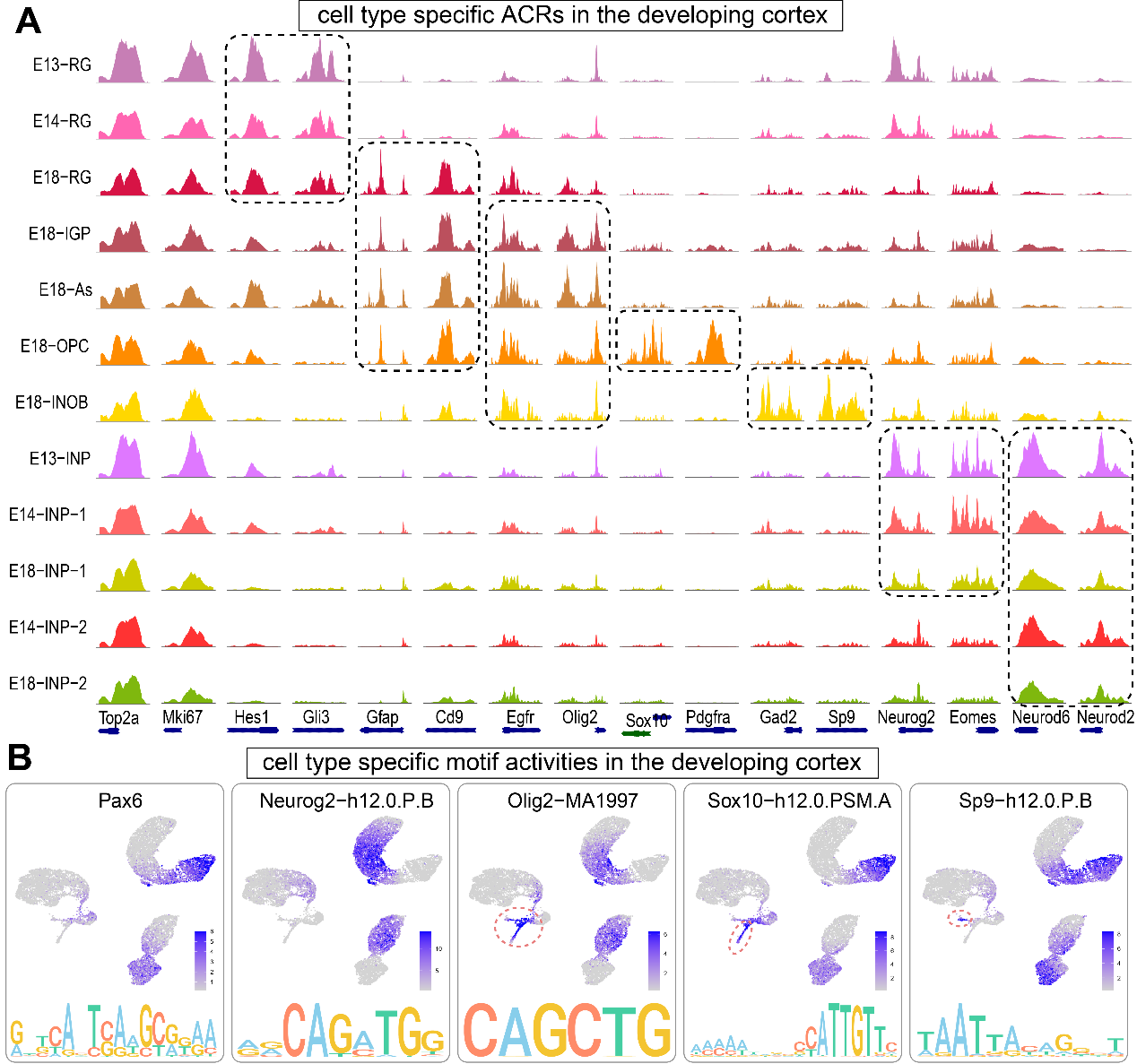

Fig. S5: **Cell-Type-Specific Chromatin Accessibility and Transcription Factor Motif Activities.** **A**: snATAC-seq coverage plots showing cell type specific chromatin accessibility during embryonic cortical development, colored by cell types. **B**: UMAP visualization of transcription factor motif activities. The circled areas indicate specific motif activities in the corresponding cell type.
